# Crossmodal reorganization in deaf auditory cortices compensates for the impaired body-centered visuomotor transformation after early deafness

**DOI:** 10.1101/2022.07.14.500143

**Authors:** Li Song, Pengfei Wang, Hui Li, Peter H. Weiss, Gereon R. Fink, Xiaolin Zhou, Qi Chen

**Affiliations:** Key Laboratory of Brain, Cognition and Education Sciences (South China Normal University), Ministry of Education, China; School of Psychology, Center for Studies of Psychological Application, and Guangdong Key Laboratory of Mental Health and Cognitive Science, South China Normal University, Guangzhou 510631, China; Cognitive Neuroscience, Institute of Neuroscience and Medicine (INM-3), Research Centre Jülich, Germany, Wilhelm-Johnen-Strasse, 52428 Jülich, Germany; Department of Neurology, University Hospital Cologne, Cologne University, 509737 Cologne, Germany; Shanghai Key Laboratory of Mental Health and Psychological Crisis Intervention, School of Psychology and Cognitive Science, East China Normal University, 200062 Shanghai, China

**Author notes:** the authors contributed equally to the present work (Li Song, Pengfei Wang). Address correspondence to: Qi Chen, PhD, or to: Xiaolin Zhou, PhD.

**Keywords:** early deafness, superior temporal gyrus, neural plasticity, frontoparietal network, default-mode network, dorsal attention network

## Abstract

Early deafness leads to the reorganization of large-scale brain networks, involving and extending beyond the auditory system. Body-centered visuomotor transformation is impaired after early auditory deprivation, associated with a hyper-crosstalk between the task-critical frontoparietal network (FPN) and the default-mode network (DMN). It remains to be elucidated, how the reorganized functional connectivity between the auditory system, the FPN, and the DMN contributes to the impaired visuomotor transformation after early deafness. In this fMRI study, we asked early deaf participants and hearing controls to judge the spatial location of a visual target, either about the middle-sagittal line of their own body (the egocentric task) or another background object (the allocentric task). The bilateral superior temporal gyrus (STG) in the deaf group exhibited cross-modal reorganization, with generally enhanced neural activity during the visual tasks, compared to hearing controls. Moreover, the STG showed significantly increased functional connectivity with both the FPN and the DMN in the deaf group compared to hearing controls, specifically during the egocentric task. The increased STG-FPN and STG-DMN coupling, however, showed antagonistic effects on the egocentric performance of the deaf participants. The increased STG-FPN connectivity was associated with improved (i.e., a beneficial role) while the increased STG-DMN with deteriorated (i.e., a detrimental role) egocentric performance in the deaf participants. No such effect was observed in hearing controls. Therefore, the auditory cortex is reorganized to functionally resemble the FPN in the deaf brain, representing compensatory neuroplasticity to mitigate the impaired visuomotor transformation after early deafness.

**Significance Statement:** Our brain constantly plans vision-guided actions, transforming visuospatial representations of external visual targets into visuomotor representations. The frontoparietal network (FPN) critically supports this visuomotor transformation process, which is impaired after early deafness. To mitigate the impaired visuomotor transformation, the ‘deaf’ auditory cortex in the bilateral superior temporal gyrus (STG) shows compensatory cross-modal reorganization that functionally resembles the FPN regions. Specifically, the deaf auditory cortex becomes functionally coupled with the dorsal FPN regions. The stronger the STG-FPN coupling, the more improved the deaf adults’ visuomotor transformation performance, indicating the reorganized STG as a critical node of the task-critical network. Correspondingly, increased coupling between the task-critical deaf STG and the default-mode network impairs the visuomotor transformation.

## Introduction

Early auditory deprivation leads to structural and functional reorganization of the auditory cortex (for reviews Amaral and Almeida, 2015; Cardin et al., 2020; Hribar et al., 2020; Simon et al., 2020). Structurally, decreased fractional anisotropy of white matter fibers, decreased white matter volume, and increased cortical thickness have been observed in the ‘deaf’ auditory cortex compared to hearing controls (Emmorey et al., 2003; Smith et al., 2011; Karns et al., 2016; Kumar and Mishra, 2018). Functionally, the ‘deaf’ auditory cortex undergoes cross-modal reorganization and starts to process stimuli from the remaining sensory modalities, such as visual and vibrotactile stimuli (Levänen et al., 1998; Karns et al., 2012; Ding et al., 2015; Bola et al., 2017; Cardin et al., 2018). Moreover, increased task-evoked or intrinsic functional connectivity between the ‘deaf’ auditory cortex and other brain regions has been revealed during a variety of cognitive tasks (Shiell et al., 2015; Ding et al., 2016; Benetti et al., 2017, 2021; Bola et al., 2017), indicating large-scale network reorganization in deaf people.

The spatial location of an object can be represented in either egocentric (i.e., relative to the viewer’s body/body effectors) or allocentric (i.e., relative to other external objects) reference frames (Paillard, 1991; Blouin et al., 1993; Burgess, 2006). The egocentric reference frames are particularly critical for guiding smooth visually-guided actions, which requires transforming visuospatial representations of external visual objects into visuomotor representations (Galati et al., 2001; Cohen and Andersen, 2002). At the neural level, the dorsal attention network (DAN) is commonly involved in coding the general visuospatial representations underlying both the egocentric and allocentric reference frames (Committeri et al., 2004; Chen et al., 2012, 2014; Gomez et al., 2014), while the frontoparietal network (FPN) is specifically involved in body-centered visuomotor transformation during the egocentric task (Galati et al., 2000; Neggers et al., 2006; Chen et al., 2012, 2014). Previous evidence consistently shows that the egocentric reference frame is impaired after early auditory deprivation (Zhang et al., 2014), associated with a hyper-crosstalk between the FPN and the default-mode network (DMN) during body-centered egocentric judgments in the deaf brain (Li et al., 2022). The DMN is generally deactivated during various externally directed tasks to suppress task-irrelevant distractions and achieve successful task performance (Anticevic et al., 2012). Accordingly, the increased connectivity between the task-critical neural network and the DMN impairs task performance (Weisz et al., 2014; Sadaghiani et al., 2015; Li et al., 2022).

Besides the large-scale network reorganization beyond the auditory system, the ‘deaf’ auditory system exhibits stronger intrinsic connections with sub-regions of the DMN and the FPN compared to hearing controls (Ding et al., 2016; Cardin et al., 2018; Andin and Holmer, 2022). It remains unclear, however, how the decreased network segregation between the auditory system, the task-critical FPN, and the DMN contributes to the impaired visuomotor transformation after early deafness. In the present fMRI study, via graph-based modularity analyses, we investigated the cross-modal reorganization of the between-module connectivity between the superior temporal gyrus (STG), the task-critical DAN and FPN, and the DMN in the deaf brain during an egocentric judgment task. If the cross-modal reorganization in the deaf STG compensates for the impaired visuomotor transformation after early deafness, we predict that: (1) the deaf STG should show enhanced between-module connectivity with the task-critical DAN and FPN, specifically during the egocentric task, compared to the hearing controls; and more importantly (2) the potentially enhanced STG-frontoparietal connectivity should be associated with improved egocentric performance in the deaf people. Moreover, suppose the deaf STG is reorganized to functionally resemble the frontoparietal regions underlying the egocentric task since the impaired visuomotor transformation after early deafness is associated with abnormal hyper-connectivity between the task-critical frontoparietal regions and the DMN (Li et al., 2022). In that case, the deaf STG should also show increased connectivity with the DMN. However, the potentially increased STG-DMN connectivity should be detrimental, rather than beneficial, to the egocentric performance of deaf people.

## Materials and Methods

### Participants

Twenty-six right-handed early deaf individuals (12 males; 21.54 ± 2.06 years old, mean ± SD) and 24 right-handed demographic-matched hearing controls (12 males; 21.58 ± 1.69 years old, mean ± SD) participated in this study. According to their self-reports, none of the deaf and normal-hearing participants had ever been clinically diagnosed with any balance problems (vestibular dysfunction). All deaf participants had congenital, profound bilateral hearing loss (> 90 dB, each ear), as determined by a standard pure-tone audiometry procedure at 500, 1000, 2000, and 4000 Hz. Hearing loss was due to genetic or pregnancy-related factors, like hereditary deafness or drug side effects. The deaf participants exhibited inconsistent speech comprehension, ranging from poor to good even with the hearing aids. Each of them was proficient in Chinese Sign Language but had poor speech articulation. The hearing participants were native Chinese speakers who had no prior hearing problems. All participants had a normal or corrected-to-normal vision, no color vision impairment, and no psychiatric or neurological diseases. More information about both groups was provided in our previous study (Li et al., 2022). Each participant had signed informed consent following the Helsinki Declaration before the experiment and got paid afterward. The Ethics Committee of the Department of Psychology, South China Normal University approved this research.

### Experimental Design

The experimental procedure was controlled using the Presentation software (Neurobehavioral Systems, RRID: SCR_002521, https://www.neurobs.com/). The stimuli consisted of a fork lying on an orange plate displayed on a gray background (Fig. 1A). The luminance of the fork was either dark (RGB: 64, 64, 64) or light (RGB: 192, 192, 192) gray. The width of the fork end was 2.5° of visual angle, and the diameter of the plate was 15° of visual angle. The fork was located at four different egocentric positions relative to the midsagittal line of the observer’s own body (i.e., −2.67°, −1.7°, 1.7°, and 2.67°) and meanwhile at four different allocentric positions relative to the midsagittal line of the plate (i.e., −3.6°, −2°, 2°, and 3.6°). The two types of positions were orthogonally crossed. At each of the four egocentric locations of the fork, the background plate was moved around the fork, forming four different allocentric positions (Fig. 1A). The visual angles of the egocentric and allocentric positions of the targets were set via an initial psychophysical test using a different group of hearing individuals to balance the task difficulty between the allocentric and egocentric judgments in the hearing controls. Our previous studies had demonstrated that these selected allocentric and egocentric positions effectively balanced the task difficulty across the three experimental tasks in the normal hearing controls (Liu et al., 2017a; Li et al., 2022).

**Figure 1.**
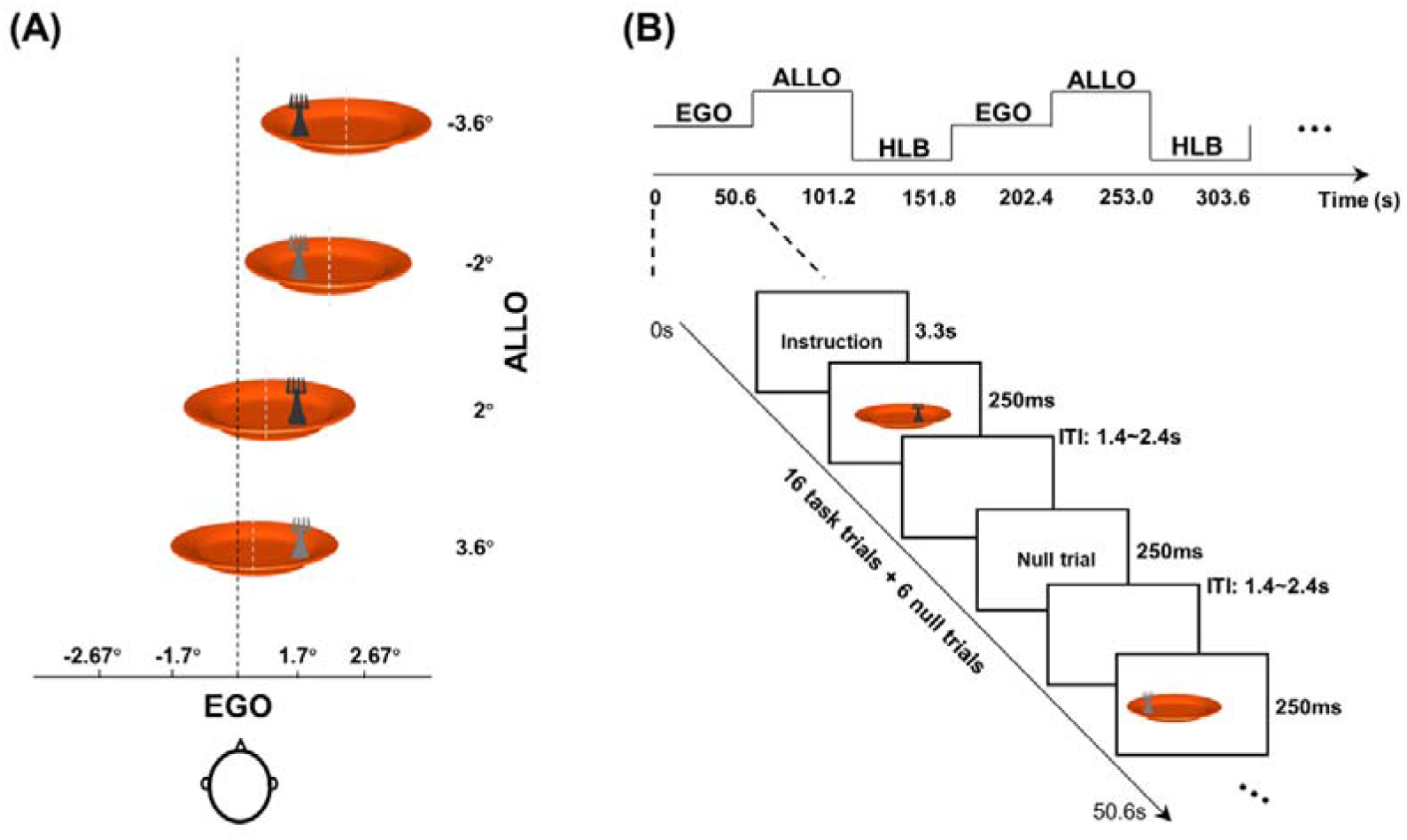
Stimuli and paradigm. (A) Stimuli. The stimuli consisted of a fork lying on a plate. The luminance of the fork was either dark or light gray. The fork was located at 4 different egocentric positions (−2.67°, −1.7°, 1.7°, and 2.67°) relative to the midsagittal line of the observer’s own body, i.e., the black vertical dashed line. Meanwhile, at each of the four egocentric locations of the fork, the background plate was moved around the fork, forming 4 different allocentric positions (−3.6°, −2°, 2°, and 3.6°) relative to the midsagittal line of the plate, i.e., the gray vertical dashed line. The egocentric and the allocentric positions were orthogonally crossed. (B) Paradigm. A mixed fMRI design was used. Three tasks were presented as alternating task blocks with pseudo-random order, and an event-related design was embedded in each task block. At the beginning of each block, a 3.3 s instruction was displayed to indicate the task of the upcoming block. In the egocentric task (EGO), participants were asked to judge the fork location relative to their own bodies’ midsagittal plane (left vs. right). In the allocentric task (ALLO), participants were asked to judge the fork location relative to the plate’s midsagittal plane (left vs. right). In the high-level baseline task (HLB), participants were asked to judge the luminance of the fork (dark vs. light grey). Within each task block, sixteen task trials and six null trials (only a blank default screen) were randomly mixed with the inter-trial intervals (ITI) jittered from 1.4 s to 2.4 s in a step of 250 ms. The target in each trial was presented for 250 ms.

All participants performed three different tasks on the same set of stimuli, including the egocentric judgment task (EGO), the allocentric judgment task (ALLO), and the non-spatial luminance discrimination task (i.e., high-level baseline, HLB). In the egocentric task, individuals judged whether the fork was on the left or right side of their bodies’ midsagittal plane. In the allocentric task, individuals judged whether the fork was on the left or right side of the plate’s midsagittal plane. For these two spatial tasks, participants had to press the left button box with their left thumb for the response of ‘left-side’ and the right button box with their right thumb for the response of ‘right-side’. In the non-spatial HLB task, individuals judged the luminance of the fork (dark gray or light gray) by pressing the left button box with their left thumb or the right button box with their right thumb. The mapping between the luminance and response hand was counterbalanced across participants. The experimental design was a 2 (between-subject factor: deaf vs. hearing) × 3 (within-subject factor: ALLO, EGO, and HLB) two-factorial design.

In this study, a mixed fMRI design was used. Three types of tasks were presented as alternating task blocks with pseudo-random order, and an event-related design was embedded in each task block (Fig. 1B). All participants alternately performed these three types of tasks 10 times without any rest block. A pseudo-random order rather than a random order of all the task blocks ensured that the maximum time interval between any two identical blocks did not exceed 200 s, thereby meeting the high-pass filter of 1/200 Hz for the following processing of the task-state fMRI data. At the beginning of each block, a 3.3 s instruction was displayed to indicate the task of the upcoming block. Within each task block, 16 task trials and 6 null trials (only a blank default screen) were randomly mixed with the inter-trial intervals jittered from 1.4 s to 2.4 s in 250 ms steps. The target in each trial was presented for 250 ms (Fig. 1B). Such a short stimulus duration was used to minimize eye movements (Findlay, 1997). The entire experiment included 160 experimental trials for each type of task and 180 null trials. Notably, no central fixation cross was presented throughout this experiment to avoid participants using it, rather than the task-required body’s midsagittal plane, as an allocentric reference object to perform the egocentric task. Participants were also asked to keep their eyes straight ahead and not move their eyeballs. In a comparable experimental paradigm, our previous research using eye-tracking technology found that central fixation could be maintained equally well in the allocentric and egocentric tasks (Chen et al., 2012). Before the formal fMRI experiment, each participant went through a training procedure to become familiar with the experimental tasks.

### Data Acquisition

The imaging data were collected using a SIEMENS 3.0T Trio Tim system with a 32-channel head coil at the Institute of Psychophysics, Chinese Academy of Sciences, Beijing. First, resting-state images were acquired using a T2-weighted EPI sequence with 200 functional volumes. The corresponding parameters as follow: slice thickness = 3 mm, repetition time = 2200 ms, echo time = 30 ms, acquisition matrix = 64 × 64, flip angle = 90°, pixel size = 3.44 × 3.44 × 3.0 mm^3^, slices = 36 with a 0.75 mm gap. This resting-state session lasted for 7.33 minutes. During this session, participants need to relax, stay awake with their eyes closed, and think about nothing. A T2-weighted EPI sequence with 729 volumes was then used to acquire individual task-state images, and the scanning parameters were the same as the resting-state session. This task-state fMRI scanning had only one run and lasted for 26.73 minutes. Finally, high-resolution structural images were acquired using a 3D MPRAGE T1-weighted sequence with 144 volumes that lasted for 8.09 minutes. The corresponding parameters were: slice thickness = 1.33 mm, repetition time = 2530 ms, echo time = 3.37 ms, inversion time = 1100 ms, acquisition matrix = 256 × 192, flip angle = *7°*, pixel size = 0.5 × 0.5 × 1.33 mm^3^.

### Analysis of Structural MRI Data

To investigate whether the anatomy of the deaf brain was altered, surface-based cortical thickness was measured in each participant, and the between-group difference was tested using the CAT12 toolbox (http://www.neuro.uni-jena.de/cat/), a well-established pipeline in SPM12 software (https://www.fil.ion.ucl.ac.uk/spm/software/spm12/) running on MATLAB R2019a. Individual structural MRI (sMRI) images were corrected for bias field and then segmented into grey matter, white matter, and cerebrospinal fluid. Subsequently, the projection-based thickness method was used to estimate the cortical thickness (Dahnke et al., 2013), and a 15mm full-width/half-maximum Gaussian kernel was used to smooth the established central surfaces. An automated quality check and further visual inspection were performed, and all structural images passed through the quality control protocol. Finally, the between-group difference in the cortical thickness was tested using the full factorial design via the CAT12. Here, a relaxed threshold with a cluster level of *p* < 0.05 (uncorrected) and a standard voxel level of *p* < 0.005 (uncorrected) was used to identify the trend of structural changes in the deaf brain since no group differences were observed at a conservative threshold of *p* < 0.05, FWE correction for multiple comparisons at the cluster level with a standard voxel level of *p* < 0.005 (uncorrected). The thickness values of regions with the structural alterations were extracted from individual sMRI data and shown as a function of the two groups.

### Network Nodes Definition

#### Preprocessing of the task-state fMRI data

Task-state fMRI data were preprocessed using SPM12 software. The preprocessing included the following steps: 1) removing the first five volumes to ensure the data were collected when the magnetic field was stable and that participants had adapted to the scanning environment, 2) slice timing to correct the phase differences, 3) realigning the functional images to the new first volume to correct head movements, 4) normalizing all images to standard MNI152 space and resampling voxel size to 3 × 3 × 3 mm^3^, and 5) smoothing with a 6mm full-width/half-maximum to alleviate the anatomical variability between participants.

#### Statistical Analysis

Preprocessed data were high-pass filtered at 1/200 Hz and modeled with a general linear model (GLM) in SPM12. The temporal autocorrelation was modeled using an AR(1) process. At the individual-level analysis, the GLM was used to construct a multiple regression design matrix. Three types of target trials (ALLO, EGO, and HLB) were modeled in an event-related analysis. The three types of neural events were time-locked to the onset of the target trials by a canonical HRF and its first-order time derivative (TD) with event duration of 0 s. Besides, the instructions, the invalid trials (missed, error, and outliers), and the six head movement parameters were modeled as another regressor of no interest. The null trials were not modeled and treated as the implicit baseline in the GLM model. Parameter estimates were calculated for each voxel using weighted least-squares to provide maximum likelihood estimators based on the temporal autocorrelation of the data. For each participant, simple main effects for each of the three experimental conditions were computed and taken to a group-level analysis. Specifically, at the group-level analysis, two factors were included in a full factorial model with one task type factor (ALLO, EGO, and HLB) and one group factor (deaf vs. hearing). To test whether the cross-modal reorganization occurred in the auditory system of the deaf participants’ brains, a *t*-test contrast of “the main effect of groups”, i.e., “Deaf (ALLO + EGO + HLB) > Hearing (ALLO + EGO + HLB)”, collapsed over all three visual tasks, was established. In this way, the reorganized regions that displayed significantly elevated neural activity during the three visual tasks in the deaf than hearing group were included. To further test whether the neural interactions between the auditory cortex and the task-related regions were altered in the deaf participants’ brains, an *F*-test contrast of “the main effect of tasks”, i.e., “ALLO vs. EGO vs. HLB”, collapsed over the two groups, was established. In this way, the brain regions that showed differential activation (positive and negative) between the three tasks were included. Areas of activation were identified as significant if they passed a threshold of *p* < 0.005, FWE correction for multiple comparisons at the cluster level with an underlying voxel level of *p* < 0.001 (uncorrected). The localized task-related regions, together with the reorganized auditory cortex, were combined to form a brain mask, and each voxel of this mask was considered a node in the subsequent modularity analyses.

### Modularity Analyses

#### Data Preprocessing

Resting-state fMRI data were preprocessed using the DPARSF module of the DPABI V6.0 software (http://rfmri.org/DPABI). The preprocessing included the following steps: 1) discarding the first five volumes, 2) slice timing correction, 3) head movement correction, 4) reorienting functional and structural images to achieve high-quality segmentation and normalization, 5) controlling for Friston-24 motion parameters, white matter signal, and cerebrospinal fluid signal as covariates, 6) normalizing functional images to standard MNI152 space using DARTEL and resampling voxel size to 3 × 3 × 3 mm^3^, and 7) bandpass filtering with 0.01-0.1 Hz. Given that removing the global signal would change the distribution of connectivity and increase negative correlations (Murphy et al., 2009; Liu et al., 2017b; Murphy and Fox, 2017), the global signal was not regressed out. The network node was each voxel in our pre-defined task mask. Hence the spatial smoothing, which would exaggerate the similarity between voxels, was not performed. Additionally, the same preprocessing of the resting-state fMRI data was also applied for the task-state fMRI data to minimize the impact of preprocessing differences and ensure the comparability of modularity results between the resting-state and the task-state.

#### Network Construction

For the resting-state data, we extracted the whole time series of each voxel from our task mask and calculated the voxel-wise Pearson correlation matrix on the individual level. For the task-state data, blocks with more than six error trials were discarded. Given the effect of hemodynamic delay, the first 4 volumes (8.8 s) were removed, and the 2 volumes (4.4 s) of the next block were included in each valid block (Mostofsky et al., 2009), so there were 20 time points per block. The time series of each voxel from our mask was extracted and a voxel-wise Pearson correlation matrix was calculated within each valid block on the individual level. All correlation matrices for each participant’s task were then averaged (Mostofsky et al., 2009; Liang et al., 2016), leaving one correlation matrix for each task type in each participant. Finally, negative associations in the correlation matrix were replaced as zero because of the controversial physiological meaning of the negative links, and a set of sparsity thresholds (2% to 5% with a step of 1%, the ratio of the number of actual edges to the maximum possible number of edges) were used to generate binary brain graphs.

#### Evaluating Network Properties

The graph-based modularity analyses were conducted on the resultant brain graphs via the GRETNA V2.0 software (https://www.nitrc.org/projects/gretna/). To identify brain modules, sets of nodes that are highly associated with each other but less associated with other modular nodes, the modified greedy optimization algorithm (Danon et al., 2006) was used in which the modularity (Q) was defined as:

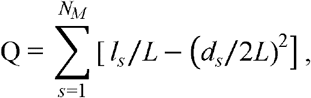

where *N_M_* is the number of non-overlapping modules, *l_s_* is the number of within-module links in the module *s, L* is the total number of links in the network, *d_s_* is the sum of degrees for each node in the module *s*, and the degree is the number of links connected to a node (Guimerà and Amaral, 2005).

For the resting-state fMRI data, we first performed the individual-level modularity analysis based on each participant’s brain graph. Since the module number and membership varied among individuals even within the same group, a group-level modularity analysis (collapsed over the two groups) was also performed to obtain a general modular structure shared by all the participants. Specifically, a group brain graph was created by averaging the correlation matrices across all participants and then thresholding at each network sparsity threshold from 2% to 5% in a step of 1%. After the group-level modularity analysis, the adjusted mutual information (AMI) was estimated to measure the similarity of module structure between the two groups. Since the modular partitions were very similar between the deaf and hearing groups based on the AMI analysis (see ‘Results’), the modules of interest comprising the DAN, the FPN, the DMN, and the bilateral superior temporal gyrus (STG) were selected from the group-level modularity analysis at the moderated network sparsity of 3%. Specifically, the DAN, the FPN, and the DMN were visually identified based on the Yeo-7 networks (Yeo et al., 2011), and the STG was identified via the Automated Anatomical Labeling (AAL) template (Tzourio-Mazoyer et al., 2002). The network properties at the modular and the nodal levels were then calculated. For the task-state fMRI data, the network properties were also estimated according to the identified modular partitions in the resting-state.

At the modular level, we evaluated the between-module connectivity for each participant. Specifically, the between-module connectivity was computed as the total number of positive links between any pair of two modules. The participation coefficient (PC) across the four modules of interest (DMN, DAN, FPN, and STG) was calculated at the nodal level. Since we were interested in the alterations of the inter-network connectivity between the STG and each of the three task-related networks (DAN, FPN, and DMN), the PC was estimated only between any pair of two modules. The PC reflects the ability of a node *i* to keep communication between its own module and the other modules, defined as:

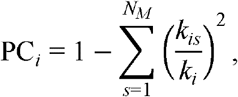

where *N_M_* is the number of non-overlapping modules (i.e., ‘2’ for the current analysis), *k_is_* is the number of positive links between the node *i* and module *s*, and *k_i_* is the sum number of positive links of node *i* in the network (including two modules in the current analysis) (Guimerà and Amaral, 2005). Therefore, if the links of node *i* are distributed across all modules, the node’s PC is close to one, and if the links of node *i* are within its corresponding module, the node’s PC is close to zero.

#### Statistical Analysis

For the module-wise analysis, we utilized the two-sample *t*-test with Bonferroni correction to examine the between-group difference in the resting-state data and the 3 (type of tasks: ALLO, EGO, and HLB) × 2 (group factor: deaf vs. hearing) ANOVA with Bonferroni correction to examine the significant interaction effect and the between-group difference in the task-state data. For the node-wise analysis, the PC image file for each participant was created based on individual nodes’ PC values, and the created images were then submitted to a group-level analysis in SPM12. More specifically, the two-sample *t*-test was implemented in the resting-state data, while the full factorial model with one task type factor (ALLO, EGO, and HLB) and one group factor (deaf vs. hearing) was implemented in the task-state data. During the task-state modularity analyses, we first examined the general changes in neural connectivity between the STG and the task-related networks (DAN, FPN, and DMN) in the deaf participants’ brains, compared to the normal hearing controls (collapsed over the three visual tasks). Moreover, we were particularly interested in the altered inter-network connectivity between the STG and the task-related networks, specifically during the egocentric task in deaf participants compared to the hearing controls. Therefore, an exclusive masking procedure was performed to eliminate the potential contribution from the other task of no interest. More concretely, the contrast “Deaf EGO > Hearing EGO” was exclusively masked by the contrast “Deaf ALLO > Hearing ALLO”, at a liberal threshold of uncorrected voxel level of *p* < 0.05. In this way, the altered neural coupling involved in the allocentric task of the deaf than the hearing group was excluded, leaving the specific alteration of neural coupling during the egocentric task in the deaf brain. The individual egocentric performance was added as a covariate in the task analysis to investigate whether and how the inter-network connections would benefit or harm the egocentric processing. Because we attempted to examine the specific effect of the egocentric performance rather than the general effect of response speed, the relative reaction time (RT) difference between the egocentric and allocentric tasks (“EGO_RT - ALLO_RT”) was used. Areas of activation were identified as significant if they passed a threshold of *p* < 0.05, FWE correction for multiple comparisons at the cluster level with an underlying voxel level of *p* < 0.05 (uncorrected). Behavior-related activation could not be corrected for multiple comparisons, but several meaningful subthreshold results were found that we believed merited further exploration. As such, an uncorrected threshold of *p* < 0.05 was used to identify a trend towards significant activation.

## Results

### Structural alternations in the deaf brain

Surface-based morphometry analyses showed increased cortical thickness in the left STG of the deaf individuals compared to the hearing controls (Fig. 2A and Table 1A).

**Figure 2.**
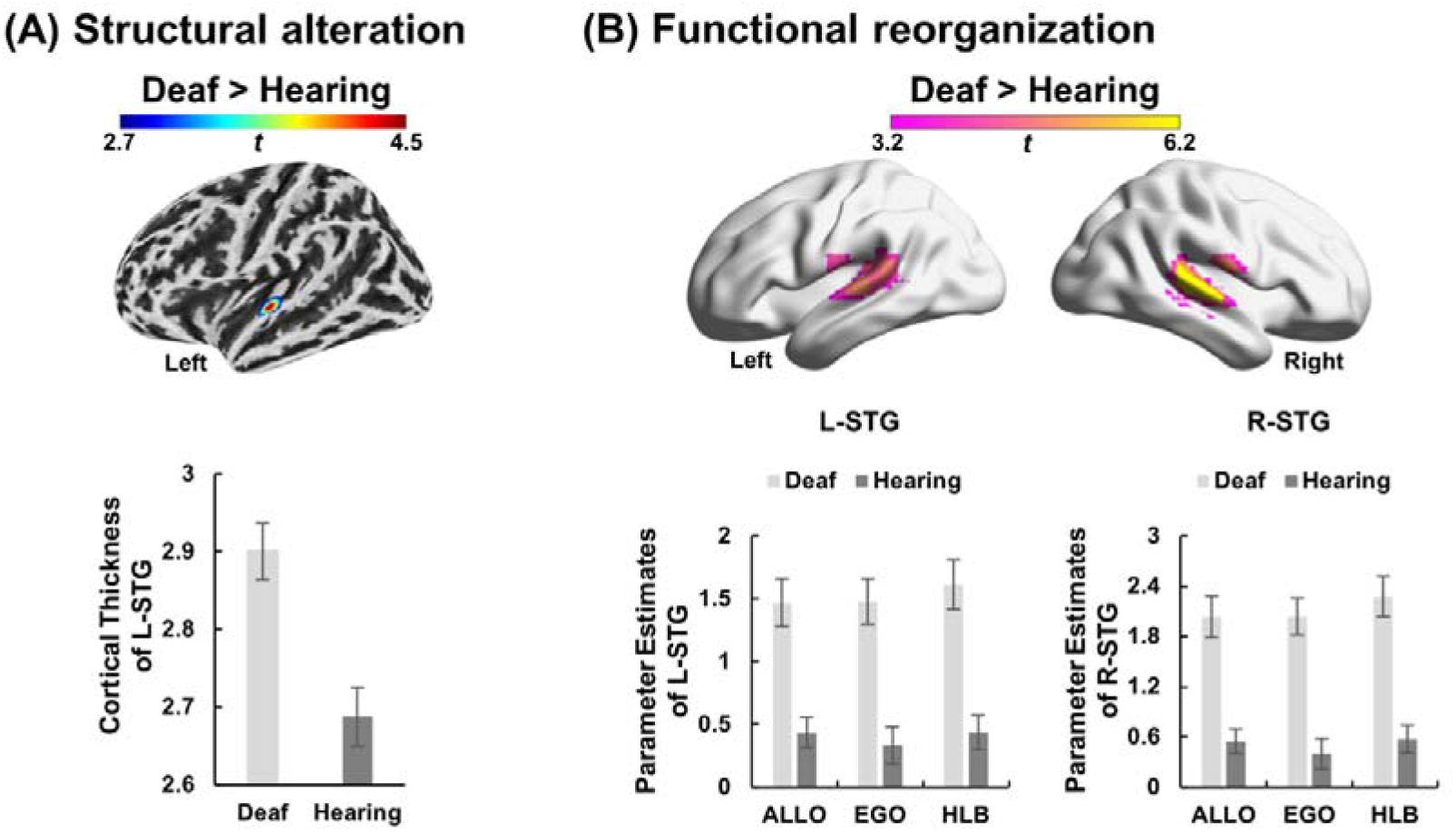
Structural alteration and functional reorganization in the deaf participants’ brains. (A) Structural alteration. The deaf group showed increased cortical thickness in the left superior temporal gyrus (STG) compared to the hearing controls. The bar plot shows the thickness values in the left STG as a function of the two subject groups. (B) Functional reorganization. Compared to the hearing controls, the deaf participants showed significantly elevated neural activity in bilateral STG in all three visual tasks (ALLO, EGO, and HLB). The bar plots show the amplitude of BOLD responses as a function of the three tasks in the two subject groups. Error bars indicate standard errors. ALLO, allocentric task; EGO, egocentric task; HLB, high-level baseline task; L, left; R, right.

**Table 1.**
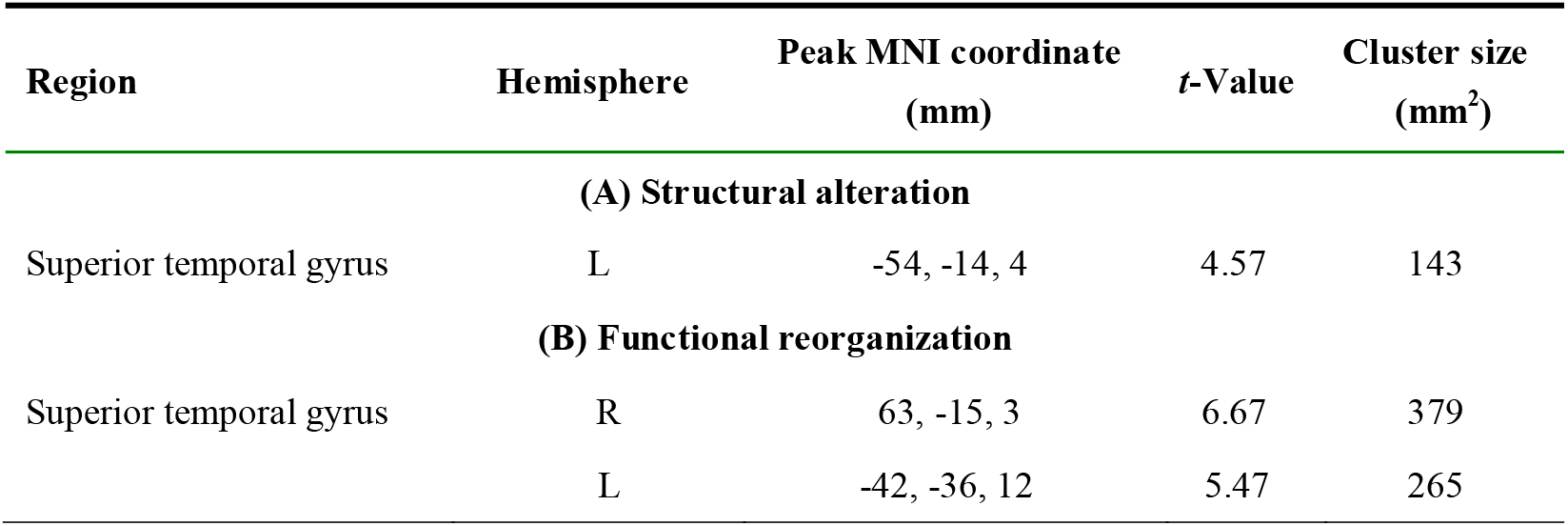
Structural alteration and functional reorganization in the superior temporal gyrus of the deaf brain.

| Region | Hemisphere | Peak MNI coordinate (mm) | *t*-Value | Cluster size (mm<sup>2</sup>) |
| --- | --- | --- | --- | --- |
| **(A) Structural alteration** |  |  |  |  |
| Superior temporal gyrus | L | -54, -14, 4 | 4.57 | 143 |
| **(B) Functional reorganization** |  |  |  |  |
| Superior temporal gyrus | R | 63, -15, 3 | 6.67 | 379 |
|  | L | -42, -36, 12 | 5.47 | 265 |

### Cross-modal responses to all the visual tasks in bilateral STG of the deaf brain

The bilateral STG was significantly activated in the neural contrast “Deaf (ALLO + EGO + HLB) > Hearing (ALLO + EGO + HLB)”, indicating that neural activity in the bilateral STG of the deaf participants’ brains generally increased upon performing the current three visual tasks, compared to the hearing controls (Fig. 2B and Table 1B).

### Brain module identification

Via an *F*-test on the task-state fMRI data, we first localized the task-related brain network involved in the differential activation (positive and negative) between any two of the three visual tasks. An extensive brain network was thus localized, including the DAN, the FPN, and the task-negative DMN (Fig. 3A). Subsequently, the localized task-related network, together with the bilateral STG activated in Fig. 2B, were combined as a brain mask image for the subsequent modularity analyses on the resting-state fMRI data.

**Figure 3.**
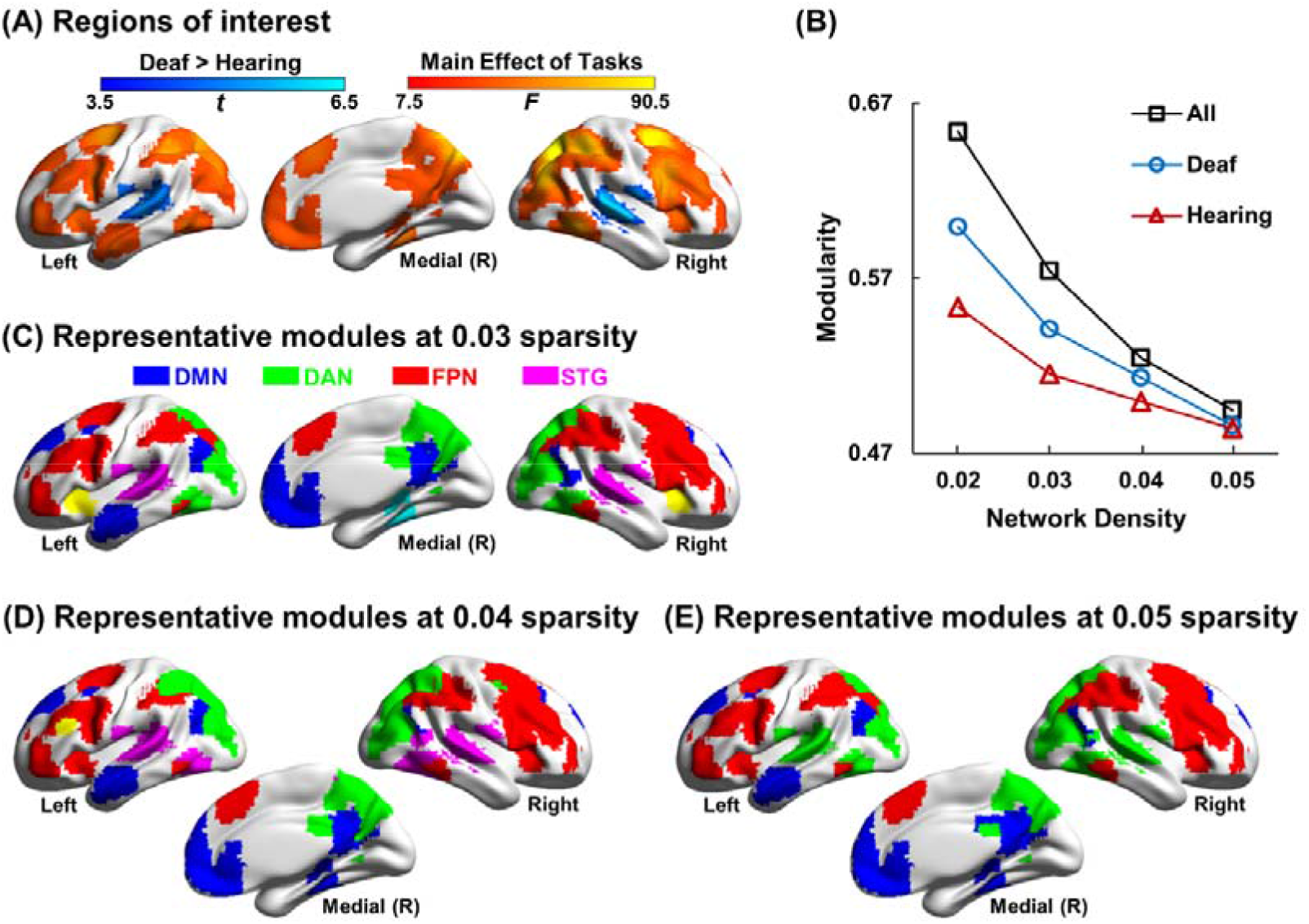
Module identification. (A) The brain mask used for module identification. The regions in warm colors are the brain regions that show differential activation (positive or negative) between any two of the three tasks “ALLO vs. EGO vs. HLB”. The regions in cold colors are the bilateral superior temporal gyrus (STG) that showed significantly elevated neural activity during the three visual tasks in the deaf than hearing group, i.e., “Deaf (EGO + ALLO + HLB) > Hearing (EGO + ALLO + HLB)”. (B) Mean modularity was obtained for the deaf group, the hearing group, and all the participants (collapsed over the two groups), respectively, with the network sparsity from 2% to 5% based on the resting-state fMRI data. (C) Map of representative modules from the resting-state data of all participants at the moderate network sparsity of 3%. The default-mode network (DMN), the dorsal attention network (DAN), the frontoparietal network (FPN), and the STG were well separated. In addition, the module structures at the sparsity threshold of 4% (D) and 5% (E) were also shown, respectively. ALLO, allocentric task; EGO, egocentric task; HLB, high-level baseline task; R, right.

Results of the modularity analyses on the resting-state data showed high modularity Q values across all the density levels, either in each participant group or in all participants combined (all more than 0.3) (Fig. 3B), indicating that the modular structure was non-random community (Newman and Girvan, 2004). Furthermore, at each network sparsity threshold, the AMI (deaf vs. hearing) ranged from 0.61 to 0.77, suggesting that the modular structure was stable and similar between the two groups (Vinh et al., 2010). Due to the similar module assignments between the two groups, the module partitions based on the group-level brain graph (collapsed over the two groups) were selected at the moderate network sparsity of 3% (Fig. 3C). Ten modules in total were identified, and four of them (STG, DAN, FPN, and DMN) were selected for the subsequent graph-based analyses on both the resting-state and the task-state data. Besides, the module structures (collapsed over the two groups) at the other two sparsity thresholds (4% and 5%) are shown in Fig. 3D and 3E, respectively.

### The resting-state: increased module connectivity between the STG, the DAN, and the FPN in the deaf brain

For the resting-state data, the between-group difference in the pair-wise connectivity between the STG and the three task-related networks (the DAN, the FPN, and the DMN) was calculated at both the modular level and the nodal level.

At the modular level, the two-sample *t*-test showed that the number of between-module connections between the STG and the DAN was significantly higher in the deaf group than the hearing group, *t*_(48)_ = 3.62, *p* = 0.001, Cohen’s *d* = 1.03 (Fig. 4A). At the nodal level, stronger inter-network connectivity between the STG and the DAN was observed in the deaf compared to the hearing group (Fig. 4B). Specifically, the right middle occipital gyrus (MOG) extending to the bilateral superior parietal lobe (SPL) and the precuneus within the DAN exhibited significantly higher PC values with the bilateral STG in the deaf than hearing group (Fig. 4B, left panel; and Table 2A). Moreover, the bilateral STG exhibited significantly higher PC values with the DAN in the deaf than hearing group (Fig. 4B, right panel; and Table 2A). Besides the DAN, the right inferior frontal gyrus (IFG) in the FPN showed significantly higher PC values with the STG in the deaf than hearing group (Fig. 4C and Table 2B). Taken together, the deaf STG showed increased connectivity with sub-regions both in the DAN and the FPN during the resting-state.

**Figure 4.**
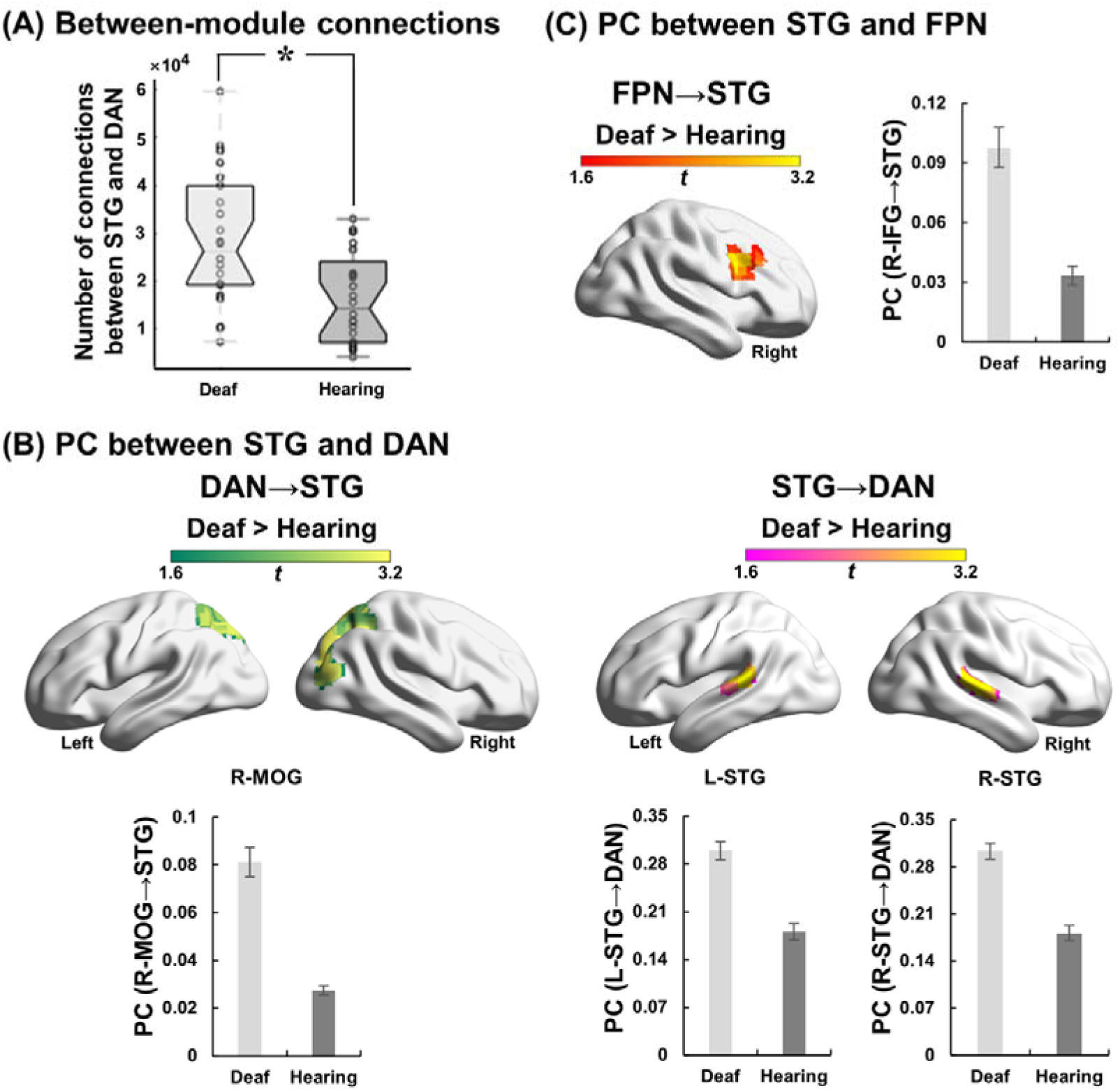
Resting-state results of module connectivity between the superior temporal gyrus (STG), the dorsal attention network (DAN), and the frontoparietal network (FPN). (A) The number of between-module connections between the STG and the DAN was significantly higher in the deaf than hearing group. Each circle represents one participant. *: *p* < 0.005. (B) Participation coefficient (PC) between the STG and the DAN. *Left panel:* the right middle occipital gyrus (MOG) extending to the bilateral superior parietal lobe (SPL) and the precuneus within the DAN exhibited significantly higher PC values with the STG in the deaf than hearing group during the resting-state. *Right panel:* the bilateral STG exhibited significantly higher PC values with the DAN in the deaf than hearing group during the resting-state. (C) The right inferior frontal gyrus (IFG) in the FPN exhibited significantly higher PC values with the STG in the deaf than hearing group during the resting-state. For demonstration purposes, mean PC values were extracted from the significantly activated regions and shown as a function of the subject group. No further statistical analysis was performed on the extracted PC values, and the error bars indicate standard errors. L, left; R, right.

**Table 2.** Participation coefficient (PC) results between the STG, the DAN, and the FPN during the resting-state.

| Region | Hemisphere | Peak MNI coordinate<br>(mm) | *t*-Value | Cluster size<br>(mm<sup>2</sup>) |
| --- | --- | --- | --- | --- |
| **(A) PC between STG and DAN** |  |  |  |  |
| **DAN→STG (Deaf > Hearing)** |  |  |  |  |
| Middle occipital gyrus | R | 30, -78, 24 | 4.81 | 449 |
| *Precuneus* | *R* | *21, -63, 24* | *4.12* |  |
| *Cuneus* | *L* | *-12, -75, 39* | *3.93* |  |
| *Superior parietal lobe* | *R* | *21, -60, 69* | *3.73* |  |
| **STG→DAN (Deaf > Hearing)** |  |  |  |  |
| Superior temporal gyrus | R | 57, -24, 9 | 5.03 | 60 |
|  | L | -54, -30, 9 | 3.50 | 42 |
| **(B) PC between STG and FPN** |  |  |  |  |
| **FPN→STG (Deaf > Hearing)** |  |  |  |  |
| Inferior frontal gyrus | R | 51, 15, 30 | 3.28 | 80 |

Italics indicate the coordinates of relevant local maxima within each significant cluster. STG, superior temporal gyrus; DAN, dorsal attention network; FPN, frontoparietal network; L, left; R, right.

### The task-state: enhanced module connectivity between the STG and the task-related networks during the egocentric task in the deaf brain

For the task-state data, at both the modular and the nodal levels, we aimed to investigate the between-group difference (deaf vs. hearing) in the inter-network connectivity between the STG and the task-related networks (i.e., DAN, FPN, and DMN), especially during the egocentric task.

At the modular level, no significant between-group difference was revealed. At the nodal level, the main effect of the subject group (‘deaf vs. hearing’, collapsed over the three visual tasks) showed that the bilateral STG of the deaf people exhibited stronger neural coupling with sub-regions in both the DAN and the FPN during all three visual tasks compared to the hearing controls (Figs. 5 and 6). More concretely, the precuneus in the DAN and the extensive regions in the FPN, including the right superior frontal gyrus (SFG), the right IFG, the right inferior parietal lobe (IPL), and the left middle frontal gyrus (MFG), showed significantly higher PC values with the STG in the deaf than hearing group upon performing the three visual tasks (Figs. 5A and 6A, the left panel; and Table 3A-B). Meanwhile, the bilateral STG showed significantly larger PC values with both the DAN and the FPN in the deaf than hearing group when performing the three visual tasks as well (Figs. 5B and 6B, the left panel; and Table 3A-B).

**Table 3.**
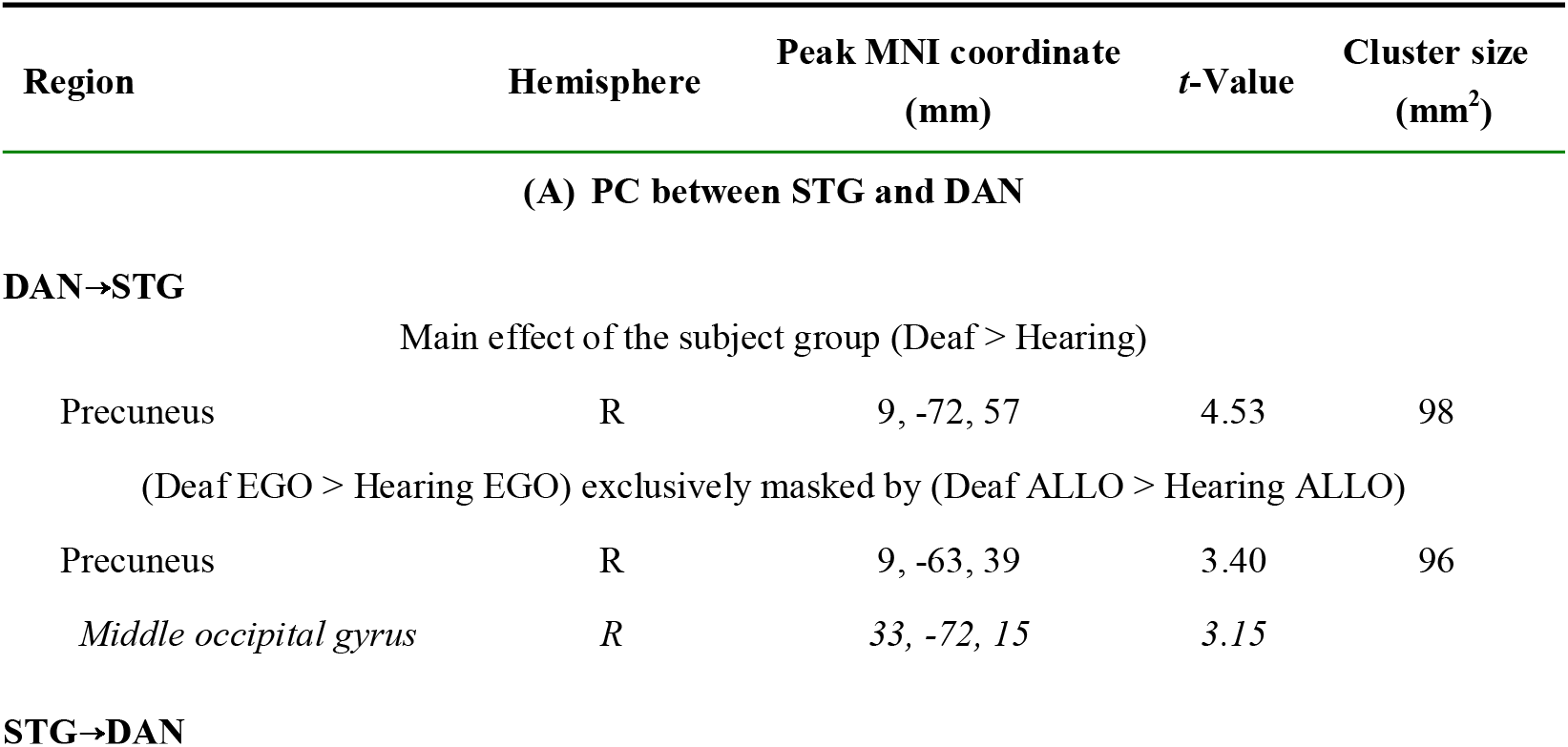
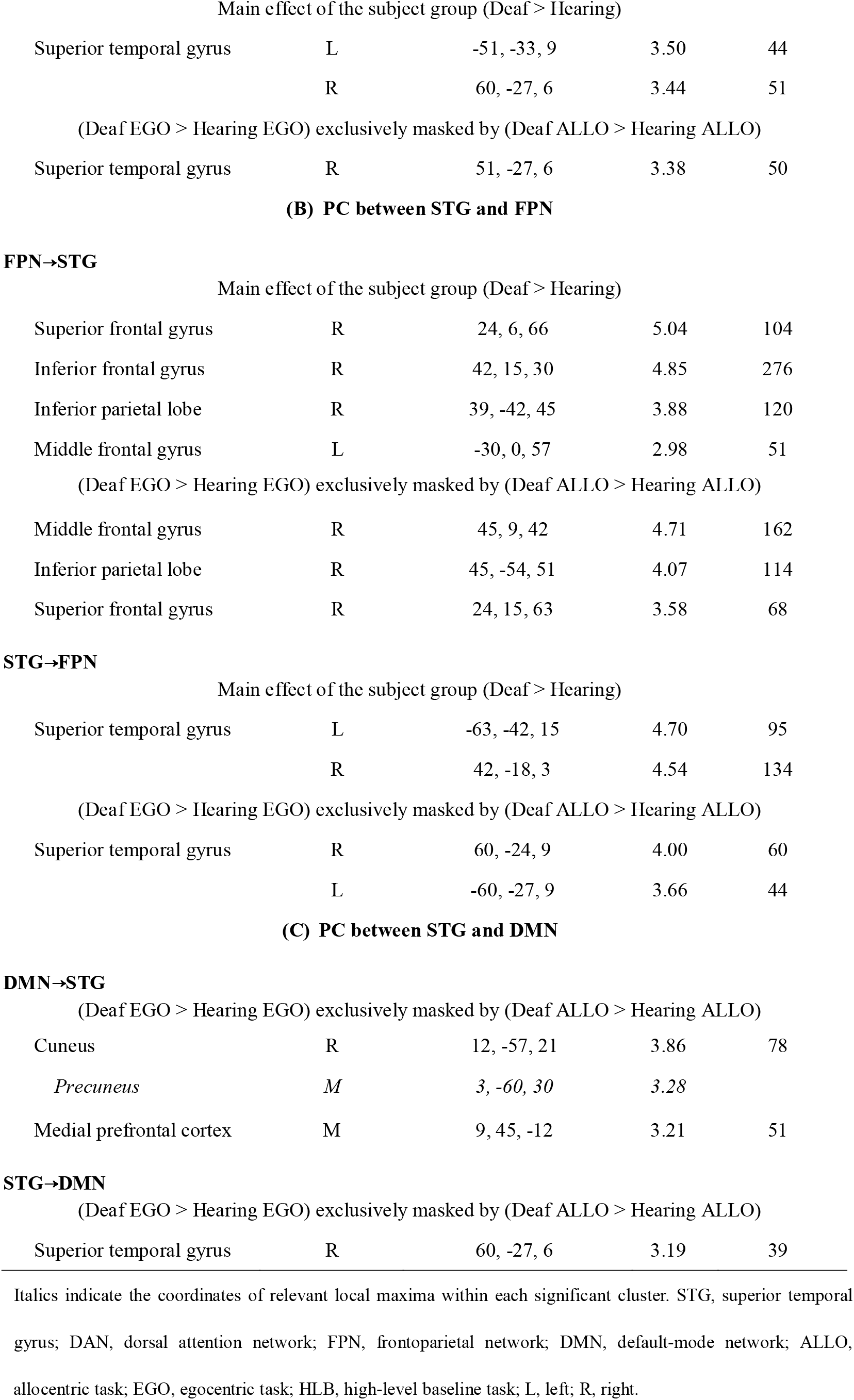
Participation coefficient (PC) results between the STG, the DAN, the FPN, and the DMN during the task-state.

| Region | Hemisphere | Peak MNI coordinate<br>(mm) | *t*-Value | Cluster size<br>(mm<sup>2</sup>) |
| --- | --- | --- | --- | --- |
| (A) PC between STG and DAN |  |  |  |  |
| **DAN→STG** |  |  |  |  |
| Main effect of the subject group (Deaf > Hearing) |  |  |  |  |
| Precuneus | R | 9, -72, 57 | 4.53 | 98 |
| (Deaf EGO > Hearing EGO) exclusively masked by (Deaf ALLO > Hearing ALLO) |  |  |  |  |
| Precuneus | R | 9, -63, 39 | 3.40 | 96 |
| *Middle occipital gyrus* | R | 33, -72, 15 | 3.15 |  |
| **STG→DAN** |  |  |  |  |

| Main effect of the subject group (Deaf > Hearing) |  |  |  |  |
| --- | --- | --- | --- | --- |
| Superior temporal gyrus | L | -51, -33, 9 | 3.50 | 44 |
|  | R | 60, -27, 6 | 3.44 | 51 |
| (Deaf EGO > Hearing EGO) exclusively masked by (Deaf ALLO > Hearing ALLO) |  |  |  |  |
| Superior temporal gyrus | R | 51, -27, 6 | 3.38 | 50 |
| **(B) PC between STG and FPN** |  |  |  |  |

FPN→STG
| Main effect of the subject group (Deaf > Hearing) |  |  |  |  |
| --- | --- | --- | --- | --- |
| Superior frontal gyrus | R | 24, 6, 66 | 5.04 | 104 |
| Inferior frontal gyrus | R | 42, 15, 30 | 4.85 | 276 |
| Inferior parietal lobe | R | 39, -42, 45 | 3.88 | 120 |
| Middle frontal gyrus | L | -30, 0, 57 | 2.98 | 51 |
| (Deaf EGO > Hearing EGO) exclusively masked by (Deaf ALLO > Hearing ALLO) |  |  |  |  |
| Middle frontal gyrus | R | 45, 9, 42 | 4.71 | 162 |
| Inferior parietal lobe | R | 45, -54, 51 | 4.07 | 114 |
| Superior frontal gyrus | R | 24, 15, 63 | 3.58 | 68 |

STG→FPN
| Main effect of the subject group (Deaf > Hearing) |  |  |  |  |
| --- | --- | --- | --- | --- |
| Superior temporal gyrus | L | -63, -42, 15 | 4.70 | 95 |
|  | R | 42, -18, 3 | 4.54 | 134 |
| (Deaf EGO > Hearing EGO) exclusively masked by (Deaf ALLO > Hearing ALLO) |  |  |  |  |
| Superior temporal gyrus | R | 60, -24, 9 | 4.00 | 60 |
|  | L | -60, -27, 9 | 3.66 | 44 |
| **(C) PC between STG and DMN** |  |  |  |  |
| DMN→STG |  |  |  |  |
| (Deaf EGO > Hearing EGO) exclusively masked by (Deaf ALLO > Hearing ALLO) |  |  |  |  |
| Cuneus | R | 12, -57, 21 | 3.86 | 78 |
| *Precuneus* | *M* | 3, -60, 30 | 3.28 |  |
| Medial prefrontal cortex | M | 9, 45, -12 | 3.21 | 51 |
| STG→DMN |  |  |  |  |
| (Deaf EGO > Hearing EGO) exclusively masked by (Deaf ALLO > Hearing ALLO) |  |  |  |  |
| Superior temporal gyrus | R | 60, -27, 6 | 3.19 | 39 |

Italics indicate the coordinates of relevant local maxima within each significant cluster. STG, superior temporal gyrus; DAN, dorsal attention network; FPN, frontoparietal network; DMN, default-mode network; ALLO,
allocentric task; EGO, egocentric task; HLB, high-level baseline task; L, left; R, right.

**Figure 5.**
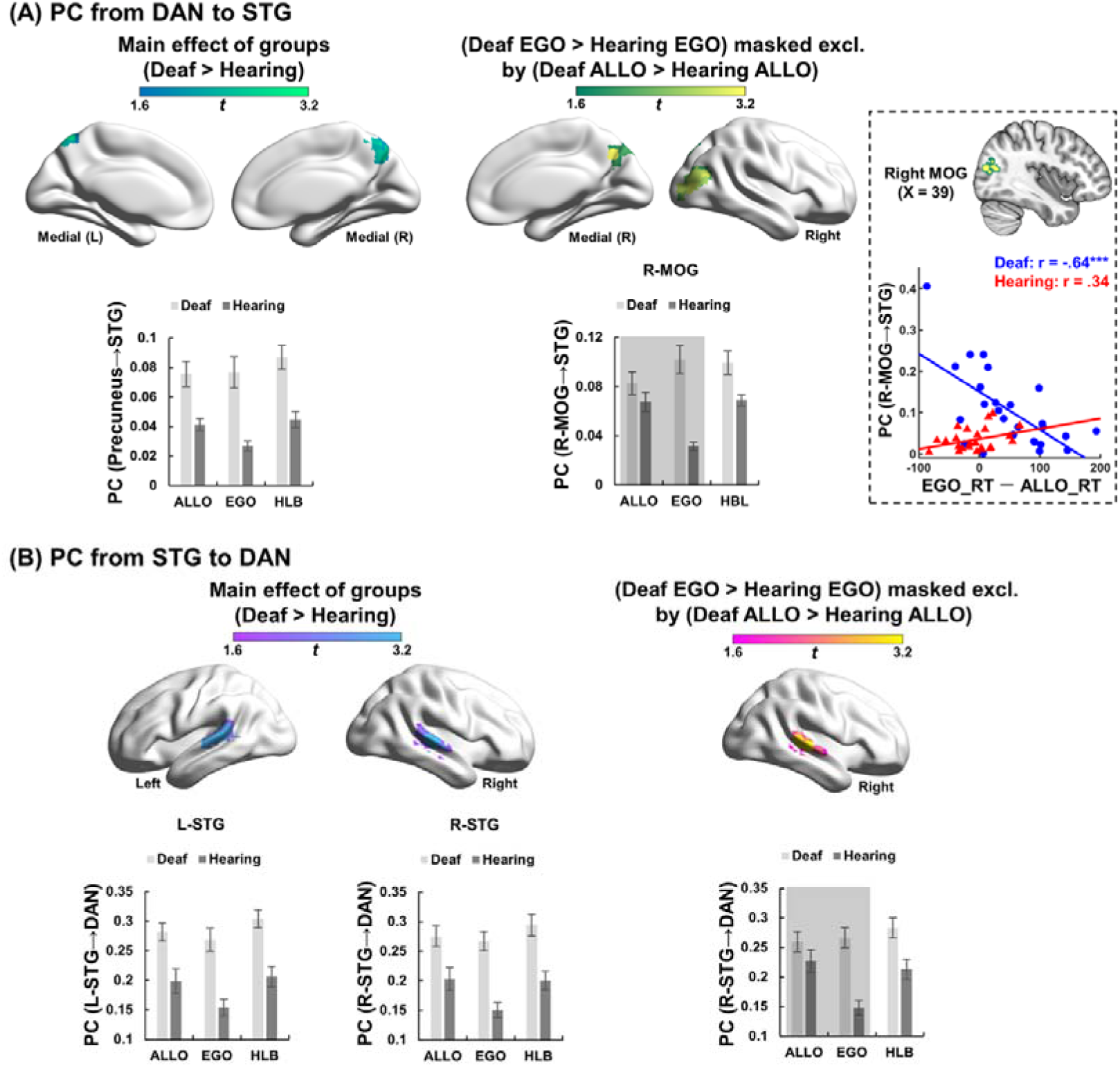
Task-state participation coefficient (PC) results between the superior temporal gyrus (STG) and the dorsal attention network (DAN). (A) PC from the DAN to the STG. *Left panel:* the main effect of subject group ‘Deaf > Hearing’ (collapsed over the three tasks). The precuneus in the DAN exhibited significantly higher PC values with the STG in the deaf than hearing group during the three visual tasks. *Right panel:* the interaction effect between the subject group (deaf vs. hearing) and the visual tasks (EGO vs. ALLO). The right middle occipital gyrus (MOG) extending to the precuneus within the DAN exhibited significantly higher PC values with the STG in the deaf than hearing group especially during the egocentric task rather than the allocentric task. Moreover, the PC value from the right MOG in the DAN to the STG was significantly negatively correlated with the egocentric performance (‘EGO_RT - ALLO_RT’) only in the deaf group but not in the hearing group. The stronger the right MOG-STG connectivity in a deaf individual, the faster the egocentric judgment. (B) PC from the STG to the DAN. *Left panel:* the main effect of subject group ‘Deaf > Hearing’ (collapsed over the three tasks). The bilateral STG exhibited significantly higher PC values with the DAN in the deaf than hearing group during the three visual tasks. *Right panel:* the interaction effect between the subject group (deaf vs. hearing) and the visual tasks (EGO vs. ALLO). The right STG showed significantly larger PC values with the DAN in the deaf than hearing group especially during the egocentric task rather than the allocentric task. For demonstration purposes, mean PC values were extracted from the significantly activated regions and shown as a function of the subject group. The conditions involved in the neural contrast were shaded. No further statistical analysis was performed on the extracted PC values, and the error bars indicate standard errors. ALLO, allocentric task; EGO, egocentric task; HLB, high-level baseline task; L, left; R, right; RT, reaction time.

**Figure 6.**
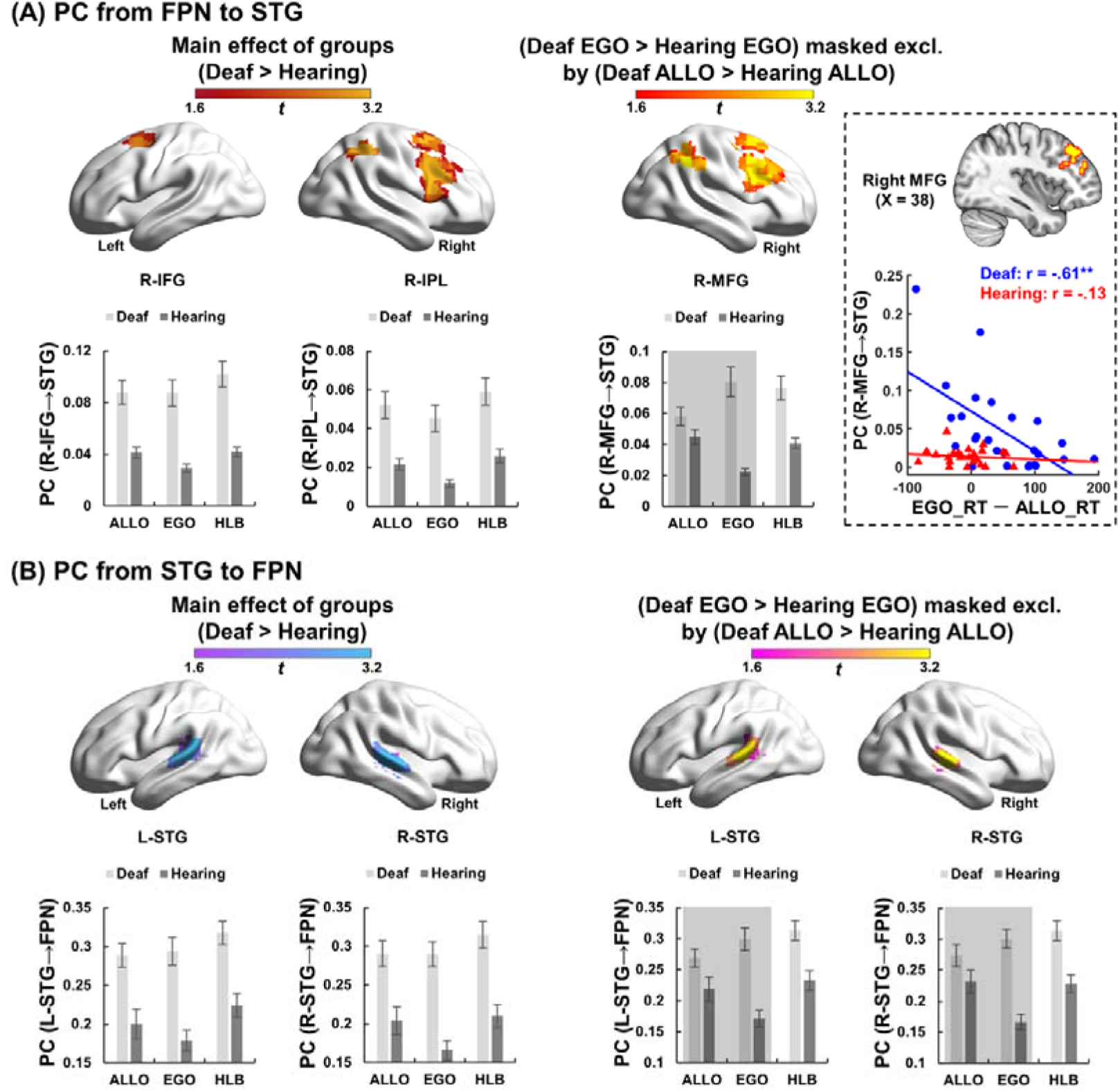
Task-state participation coefficient (PC) results between the superior temporal gyrus (STG) and the frontoparietal network (FPN). (A) PC from the FPN to the STG. *Left panel:* the main effect of subject group ‘Deaf > Hearing’ (collapsed over the three tasks). Extensive areas in the FPN exhibited significantly higher PC values with the STG in the deaf than hearing group during the three visual tasks. *Right panel:* the interaction effect between the subject group (deaf vs. hearing) and the visual tasks (EGO vs. ALLO). Extensive areas in the FPN, including the right inferior parietal lobe (IPL), the right superior frontal gyrus (SFG), and the right middle frontal gyrus (MFG), exhibited significantly stronger PC values with the STG in the deaf than hearing group especially during the egocentric task rather than the allocentric task. Moreover, the PC value from the right MFG in the FPN to the STG was significantly negatively correlated with the egocentric performance (‘EGO_RT - ALLO_RT’) only in the deaf group but not in the hearing group. The stronger the right MFG-STG connectivity in a deaf individual, the faster the egocentric judgment. (B) PC from the STG to the FPN. *Left panel:* the main effect of subject group ‘Deaf > Hearing’ (collapsed over the three tasks). The bilateral STG exhibited significantly higher PC values with the FPN in the deaf than hearing group during the three visual tasks. *Right panel:* the interaction effect between the subject group (deaf vs. hearing) and the visual tasks (EGO vs. ALLO). The bilateral STG showed significantly larger PC values with the FPN in the deaf than hearing group especially during the egocentric task rather than the allocentric task. For demonstration purposes, mean PC values were extracted from the representative significantly activated regions and shown as a function of the three tasks in each subject group. The conditions involved in the neural contrast were shaded. No further statistical analysis was performed on the extracted PC values, and the error bars indicate standard errors. ALLO, allocentric task; EGO, egocentric task; HLB, high-level baseline task; L, left; R, right; RT, reaction time; IFG, inferior frontal gyrus.

More critically, we found enhanced connectivity between the STG and sub-regions of the DAN in the deaf than hearing group, specifically during the egocentric task (Fig. 5). More concretely, within the DAN, the right MOG extending into the precuneus showed significantly higher PC values with the STG in the deaf than hearing group especially during the egocentric task, rather than the allocentric task (Fig. 5A, the right panel; and Table 3A). Moreover, the PC value from the right MOG to the STG was significantly negatively correlated with the individual difference in the egocentric performance only in the deaf group, but not in the hearing group: the stronger the right MOG-STG connectivity in a deaf individual, the faster her/his egocentric judgment (Fig. 5A, the right-most panel). On the other hand, the right STG also showed significantly larger PC values with the DAN in the deaf than hearing group, especially in the egocentric task, rather than the allocentric task (Fig. 5B, the right panel; and Table 3A).

Besides the DAN, enhanced inter-network connectivity between the STG and the FPN, specifically during the egocentric task, was also found in the deaf rather than the hearing group (Fig. 6). The right IPL, the right SFG, and the right MFG within the FPN exhibited significantly stronger PC values with the STG in the deaf than hearing group especially in the egocentric task, rather than the allocentric task (Fig. 6A, the right panel; and Table 3B). Moreover, the PC value from the right MFG to the STG was significantly negatively correlated with the individual difference in the egocentric performance only in the deaf group, but not in the hearing group: the stronger the right MFG-STG connectivity in a deaf individual, the faster her/his egocentric judgment (Fig. 6A, the right-most panel). On the other hand, the PC values from the bilateral STG to the FPN were also significantly higher in the deaf than hearing group, specifically during the egocentric task, rather than the allocentric task (Fig. 6B, the right panel; and Table 3B).

### The task-state: enhanced module connectivity between the STG and the DMN during the egocentric task in the deaf brain

For the inter-network connectivity between the STG and the DMN, no significant between-group difference (collapsed over the three visual tasks) was revealed. However, stronger inter-network connectivity between the STG and the DMN was found in the deaf than hearing group, only during the egocentric rather than allocentric task (Fig. 7). Specifically, the medial prefrontal cortex (mPFC) and the posterior cingulate cortex (PCC) in the DMN exhibited significantly larger PC values with the STG in the deaf than hearing group, especially during the egocentric rather than allocentric task (Fig. 7A, the left panel; and Table 3C). Moreover, the PC value from the PCC to the STG was significantly positively correlated with the individual difference in the egocentric performance only in the deaf group, but not in the hearing group: the stronger the PCC-STG connectivity in a deaf individual, the slower her/his egocentric judgment (Fig. 7A, the right panel). On the other hand, the right STG also showed significantly larger PC values with the DMN in the deaf than hearing group during the egocentric rather than allocentric task (Fig. 7B and Table 3C).

**Figure 7.**
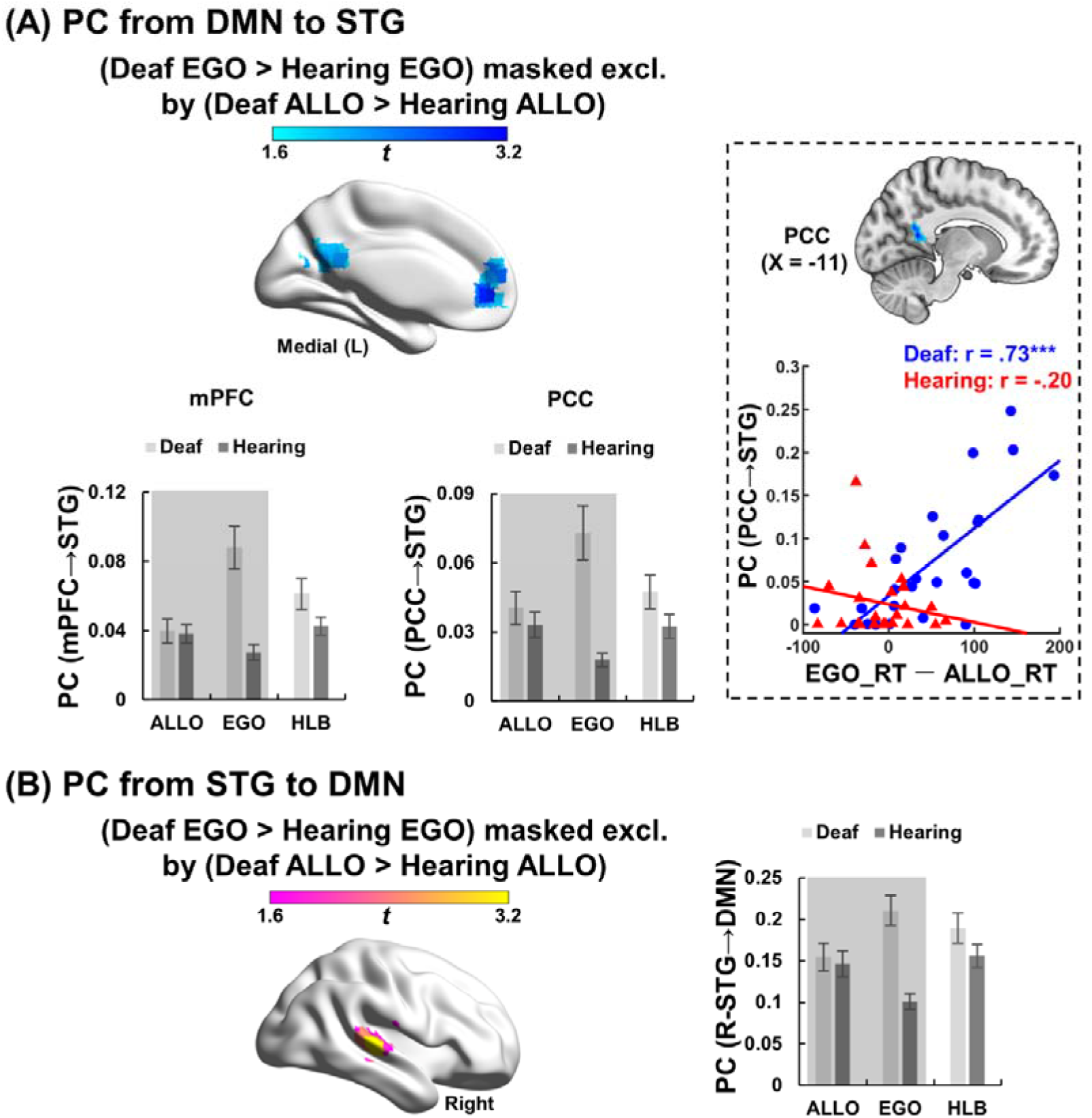
Task-state participation coefficient (PC) results between the superior temporal gyrus (STG) and the default-mode network (DMN). (A) PC from the DMN to the STG. *Left panel:* the interaction effect between the subject group (deaf vs. hearing) and the visual tasks (EGO vs. ALLO). Both the medial prefrontal cortex (mPFC) and the posterior cingulate cortex (PCC) in the DMN showed significantly stronger PC values with the STG in the deaf than hearing group especially during the egocentric rather than the allocentric task. Moreover, the PC value from the PCC in the DMN to the STG was significantly positively correlated with the egocentric performance (‘EGO_RT - ALLO_RT’) only in the deaf group, but not in the hearing group (*right panel*). The stronger the PCC-STG connectivity in a deaf individual, the slower the egocentric judgment. (B) PC from the STG to the DMN. The right STG showed significantly stronger PC values with the DMN in the deaf than hearing group during the egocentric rather than allocentric task. For demonstration purposes, mean PC values were extracted from the significantly activated regions and shown as a function of the three tasks in each subject group. The conditions involved in the neural contrast were shaded. No further statistical analysis was performed on the extracted PC values, and error bars indicate standard errors. ALLO, allocentric task; EGO, egocentric task; HLB, high-level baseline task; L, left; RT, reaction time.

## Discussion

Body-centered visuomotor transformation during egocentric judgments is impaired after early auditory deprivation. This behavioral impairment is associated with a hyper-crosstalk between the DMN and the task-critical networks in the DAN and FPN (Li et al., 2022). Here, we investigated the cross-modal reorganization in the auditory cortex of early deaf people during body-centered visuomotor transformation focusing on altered neural network dynamics between the auditory cortex in the bilateral STG, the task-critical networks in the DAN and FPN, and the DMN during the egocentric task, compared to the allocentric task, in early deaf people.

Morphologically, increased cortical thickness in the left STG of the deaf group was observed in this study (Fig. 2A). All previous surface-based morphometry studies with a relatively small sample size (less than sixteen participants per group) showed no significant difference in the cortical thickness of the auditory regions between the deaf and hearing groups (Li et al., 2012; Hribar et al., 2014; Smittenaar et al., 2016). This lack of significant differences is likely because the cortical folds in the temporal lobe are inconsistent across individuals. Thus larger sample sizes may be required to detect the temporal lobe’s thickness changes (Pardoe et al., 2012). Accordingly, two recent studies with larger sample sizes (thirty and fifty participants per group, respectively) showed that the cortical thickness of the STG increased in deaf relative to hearing people (Kumar and Mishra, 2018; McCullough and Emmorey, 2020). The initial increase in synaptic density related to synaptogenesis is not supported by the sensory input (Winfield, 1981; Bourgeois and Rakic, 1996). Instead, the subsequent synaptic pruning with the removal of unnecessary synapses and neurons, accompanied by cortical thinning, is supported by sensory experiences (Bourgeois et al., 1989; Yu et al., 2013; Faust et al., 2021). Also, the abnormal cortical thickness may be related to cortical malformation (Hyde et al., 2007; Hogstrom et al., 2012). In the tension-based theory, cortical folding can be explained by mechanical tension along axons, dendrites, and glial processes (Van Essen, 1997). Early auditory deprivation may change the tonotopic organization of the deaf brain, related to a thicker auditory cortex (for review, Hribar et al., 2020). Therefore, the increased cortical thickness of the auditory cortex in the deaf participant’s brain suggests that the lack of early auditory input results in inadequate synaptic pruning and cortical malformation. Functionally, we found that the deaf auditory cortex in bilateral STG was generally hyper-activated during all three visual tasks, as compared to the hearing controls (Fig. 2B). Early auditory deprivation alters an individual’s interaction with the external environment, which leads to a striking functional reorganization in the deaf auditory system. Accordingly, mounting empirical evidence shows that the ‘deprived’ auditory cortex of deaf people is recruited by the remaining senses, such as the visual and vibrotactile stimuli (Bavelier et al., 2000, 2001; Fine et al., 2005; Karns et al., 2012; Cardin et al., 2013, 2018; Ding et al., 2015; Benetti et al., 2017, 2021).

The functional cross-modal reorganization in the deaf brain is not only confined within the auditory system but also manifests as altered cortico-cortical connectivity between the auditory cortex and other cortical regions during a variety of visual tasks (Bavelier et al., 2000; Shiell et al., 2015; Benetti et al., 2017, 2021; Bola et al., 2017). It remains unclear, however, whether the altered neural network dynamics between the STG and other cortical areas were beneficial or detrimental to a specific visual task. In the present study, the auditory system in the bilateral STG showed enhanced functional connectivity with the DAN and the FPN in the deaf people (compared to the hearing controls) during both the resting-state (Fig. 4) and the egocentric task (Figs. 5 and 6). Moreover, the stronger the functional connectivity during the egocentric task between the STG and the DAN as well as between the STG and the FPN, the better deaf persons performed in the egocentric task (Figs. 5A and 6A, the right-most panel). This finding indicates a beneficial role of the enhanced STG-task network connectivity. Previous studies suggest that during the egocentric task, the DAN supports general visuospatial representations (Committeri et al., 2004; Chen et al., 2012, 2014; Gomez et al., 2014; Liu et al., 2017a), while the FPN supports the body-centered visuomotor transformation (Galati et al., 2000; Neggers et al., 2006; Chen et al., 2012, 2014; Liu et al., 2017a). To ensure efficient task performance, the task-relevant regions are highly connected to maintain high modularity of the task-relevant network, and meanwhile, the task-relevant regions are disconnected from the task-irrelevant regions (Ekman et al., 2012; Gonzalez-Castillo et al., 2012; Gratton et al., 2016). In the present study, the STG was not involved in the egocentric task for the hearing controls (Fig. 2B). Therefore, the connectivity between the task-irrelevant STG and the task-relevant networks (DAN and FPN) was minimized when the hearing controls performed the egocentric task. For deaf individuals, however, both the reorganized auditory system in the STG and the task-relevant DAN and FPN were involved in the egocentric task (Figs. 2B and 3A). Therefore, the neural coupling between the STG and the DAN and the FPN was enhanced to optimize the egocentric performance: the higher the neural coupling, the better the egocentric performance in the deaf people (Figs. 5A and 6A, the right-most panel). Previous studies also showed that a stronger interaction between the task-related networks benefitted cognitive task performance (Sala-Llonch et al., 2012; Thompson et al., 2013; Sadaghiani et al., 2015; Spadone et al., 2015; Shine et al., 2016). Together with previous evidence, the present results imply that the deaf STG becomes functionally analogous with the DAN and the FPN during the egocentric task. The efficient information flow between the STG and the cortical areas for egocentric task performance mitigated the impaired body-centered visuomotor transformation after early auditory deprivation.

On the other hand, the deaf auditory cortex also showed enhanced connectivity with the task-irrelevant DMN during the egocentric task (Fig. 7). In contrast to the enhanced STG-task-relevant network connectivity, the enhanced STG-task-irrelevant DMN connectivity impaired the egocentric performance of the deaf people: the stronger the STG-DMN connectivity, the worse the egocentric performance in the deaf people (Fig. 7A, the right panel). Efficient task performance was associated with stronger modularity in the task-negative DMN in terms of higher within-module connectivity in the DMN and lower between-module connectivity between the DMN and the task-relevant neural networks (Gonzalez-Castillo and Bandettini, 2017). It has been suggested that the DAN and the FPN, two task-critical networks supporting the egocentric task, show stronger functional and structural connectivity with the DMN in early deaf people (Dell Ducas et al., 2021; Li et al., 2022). Moreover, the increased inter-network connectivity between the task-irrelevant DMN and the task-relevant DAN and FPN was associated with impaired egocentric performance in congenitally deaf people (Li et al., 2022). Given that the deaf participants’ STG became functionally analogous with the task-relevant DAN and FPN during the egocentric processing (Figs. 5 and 6), the increased neural coupling between the task-relevant STG and the task-irrelevant DMN became detrimental, rather than beneficial, to the egocentric performance in the deaf people.

Since all deaf participants in the present study were sign-language users, we cannot rule out sign-language experience impacting the differences in large-scale inter-network connections between the deaf and hearing groups. Further research with native deaf signers, native hearing signers, and hearing non-signers, is warranted to tease apart the effect of early auditory deprivation vs. sign-language experience on the large-scale functional reorganization of the auditory system during visuomotor transformation.

To summarize, we revealed extensively reorganized inter-network connectivity between the auditory system in the bilateral STG and the task-relevant DAN and FPN and between the bilateral STG and the task-irrelevant DMN in the deaf brain during body-centered visuomotor transformation. The STG in deaf persons seems to become functionally analogous with the task-relevant regions in the deaf DAN and FPN during the egocentric task. Therefore, the more substantial the connectivity between the STG of deaf persons and the task-relevant networks enhanced the egocentric performance, while the stronger connectivity between their STG and the task-irrelevant DMN impaired their egocentric performance.

## Acknowledgments

The Natural Science Foundation of China grants (31871138, 32071052) supported this work.

